# Intestinal Stem Cells from Patients with Inflammatory Bowel Disease Retain an Epigenetic Memory of Inflammation

**DOI:** 10.1101/2025.05.24.655923

**Authors:** Feda H. Hamdan, Isaiah Perez, Kimberlee Kossick, Hannah Smith, Adam Edwinson, Jose M. de Hoyos-Vega, Michelle Gonzalez, David Chiang, Emily Klatt, Kristy Rumer, Mauricio Perez Pachon, Mary Sagstetter, Jessica Friton, Erin Kammer, Alana English, Lucas C.S. Chini, Jarl F. Carnahan, Noah A. Baca, Rohini Mopuri, Aditya Bhagwate, Laura E. Raffals, Zhifu Sun, Madhusudan Grover, Alexander Revzin, William A. Faubion, Brooke R. Druliner

## Abstract

Intestinal epithelial damage and impaired repair are hallmarks of ulcerative colitis (UC), even after inflammation resolves. Intestinal stem cells (ISCs) can retain stable epigenetic changes after inflammation, highlighting the potential for long-lived epithelial memory in the gut. Inflammatory injury in barrier tissues induces epigenetic memory in epithelial stem cells, and the tendency of UC to relapse at previously inflamed sites led us to hypothesize that ISCs from IBD patients acquire lasting memory of prior inflammation. To test this, we derived colonic organoids from inflamed and uninflamed regions of the same UC patients and propagated in long-term culture. Chromatin profiling revealed 2,252 accessible regions unique to prior-inflamed (PI) organoids, associated with stress response, repair, and inflammatory genes. Although these regions remained accessible, ∼95% of associated genes were not upregulated in PI organoids, indicating a primed state. Upon inflammatory or injury re-challenge, PI organoids exhibited heightened transcriptional responses and accelerated wound closure, despite reduced clonogenicity and impaired barrier function, indicating a retained inflammatory memory program. Our findings demonstrate that human ISCs retain a chromatin-based memory of inflammation that persists in the absence of immune cues and shapes future responses to injury. While this may support epithelial adaptation to secondary insults, it may predispose tissue to relapse in patients with UC.

## INTRODUCTION

Inflammatory bowel disease (IBD) comprises chronic, relapsing disorders of the gastrointestinal tract, including ulcerative colitis (UC), which is a relapsing and remitting disease with little insight into mechanisms governing perpetual cycles of inflammation and impaired mucosal healing.(1–3) Despite advanced immunotherapy, only 30-35% of patients experience substantive remission, and most patients suffer >3 relapses per year.(4–7) UC is unique in that 15-20% of patients experience limited disease with sharp demarcation between continuous inflammation and histologically normal mucosa.(8) While the pathophysiologic explanation of such sharp, limited demarcation of inflammation is unknown, this feature provides an opportunity to study healthy and diseased tissue in the same patient. Current therapies focus exclusively on suppressing immune activity, with the goal of reducing inflammation to allow the intestinal epithelium to repair. Incomplete mucosal healing persists in many patients despite remission, highlighting the need to define how chronic inflammation affects epithelial repair.(9, 10)

The intestinal epithelium is one of the most rapidly renewing tissues in the body, undergoing complete turnover every 3-5 days. This regenerative capacity is fueled by intestinal stem cells (ISCs), located at the base of the crypts, which maintain epithelial homeostasis and mount rapid repair responses following injury. In response to inflammation, infection, or physical damage, ISCs activate stress-response programs that promote proliferation and coordinate epithelial regeneration.(11–13) Although the intestinal epithelium often appears histologically normal following resolution of inflammation, IBD is marked by a relapsing-remitting disease course, suggesting that full molecular recovery may not always accompany morphological healing. This raises the possibility that inflammation leaves behind a molecular memory of that experience in ISCs that influences their future behavior.

Emerging studies in IBD have shown that the intestinal epithelium can retain stable epigenetic modifications after inflammatory exposure. In Crohn’s disease, patient-derived organoids display persistent hypomethylation at MHC class I loci, linked to altered epithelial-immune interactions. (14) In a murine model of gastrointestinal graft-versus-host disease, intestinal stem cells exhibit durable chromatin and DNA methylation changes that impair regenerative capacity.(15) These findings support the broader concept that epithelial memory may contribute to disease pathogenesis in the gut.

While memory in immune cells has been recognized for decades as central to host defense, (16–20) recent studies in murine skin have established that non-immune cells such as epithelial stem cells can also encode prior inflammatory experiences at the epigenetic level. (21) This form of long-lasting epigenetic reprogramming is called inflammatory memory, where inflammatory insults can induce lasting changes in chromatin accessibility, leaving stem cells in a primed transcriptional state. This memory is mediated by transcription factors such as AP-1 (FOS/JUN) and STAT3, which preserve accessibility at specific enhancer regions and enable rapid reactivation of target genes during subsequent injury.(22) Inflammatory memory has also been observed in other tissues, including the pancreas and developing intestine following early-life infection, suggesting it may represent a conserved epithelial response to injury.(23–25) Understanding how prior inflammation alters ISC function may help explain why epithelial repair remains incomplete in many UC patients, even during periods of clinical remission, and may offer insight into mechanisms driving relapse. The contribution of inflammatory memory to ISCs, and the consequences for the intestinal epithelium in chronic inflammatory disease such as IBD is the central aim of our study.

To investigate how inflammatory exposures affect ISC molecular and functional properties, we used intestinal organoids, three-dimensional cultures that preserve epithelial diversity and support long-term ISC expansion.(26–29) Organoids provide a powerful, human-based model to study epithelial-intrinsic biology in IBD, independent of immune signals. (30–34) Prior studies have shown that organoids can preserve transcriptional programs reflective of their tissue of origin, including disease-associated gene expression patterns.(35) While emerging evidence from cancer and developmental models suggests that organoid systems can retain epigenetic features, this has not been systematically examined in the context of IBD.(35–37) In the present study, we generated patient-matched organoids from inflamed and uninflamed colonic tissue in individuals with UC, enabling direct, intra-patient comparisons of epithelial state. By integrating chromatin and transcriptional profiling, we identify a conserved program of inflammatory memory in human ISCs that persists after inflammation resolves and alters subsequent regenerative responses of the ISC.

## METHODS

### Sex as a biological variable

Both male and female participants were included in this study. Colonic tissue biopsies were obtained from UC patients of both sexes undergoing endoscopy. The study design focused on epithelial-intrinsic mechanisms of inflammatory memory, which are not expected to be specific to one sex. Due to the paired, within-patient design, statistical comparisons of sex effects were not performed, and the study was not powered for sex-based subgroup analyses.

### Human organoid generation and culture

All patient-derived organoids used in this study were collected from Mayo Clinic patients following informed consent (please see “Study Approval” section below). Information on tissue location and macroscopic inflammation status was registered at the time of sample collection. Organoids were generated following the isolation of crypts or single cells from fresh tissues collected during biopsy or surgery using established procedures.(26, 28) Briefly, crypts were released from freshly collected tissue using 5mM EDTA, rocking at 4°C for 60-75 min. Isolated crypts were then embedded in ice-cold Matrigel (Corning® Matrigel® Growth Factor Reduced Product #356231), plated in 24-well plates and overlaid with Human Colon Media. For passaging, organoids were collected and digested in TrypLE Express (Gibco, #12604013) until small fragments or single cells were produced, which were counted. Media was changed every 3 days. See Supplementary Methods for expanded details.

### Organoid Forming Efficiency (OFE)

Organoids were dissociated into single cells using TrypLE Express (Gibco, #12604013) and were strained using a Falcon 40 uM cell strainer (Corning, #352240). The single cell suspension was counted on a hemocytometer. The single cells were plated in 8 ul domes of 100% Matrigel (Corning, #35623) at 5×10^3 cells/dome in a 96 well flat-bottom cell culture plate (Starstedt, #83.3924.500). After plating, 100 ul of Human Colon Media was added to each well, and 50 ul of Human Colon Media was added to each well every 2-3 days. On day 7 post-passage, the whole domes were imaged in z-stacks at 2.5x on a Zeiss LSM780 confocal microscope in the two-photon microscopy transmission (TPMT) setting. Organoids were identified and counted in Fiji/ImageJ manually using the cell counter tool. Organoid forming efficiency was calculated by dividing the number of organoids formed day 7 post-passage by the total number of cells seeded. Analysis and calculations were performed in GraphPad Prism using a parametric t-test.

### Assessment of barrier function in NI and PI organoid monolayers

For polarization, organoid fragments were seeded onto the apical chamber of 12-well Transwells (6 mm diameter; 3 μm pore size, Corning, US); See Supplementary Methods for seeding organoids as monolayers on transwells. Culture media was replaced every 3 days. Initially, transepithelial electric resistance (TER) was measured manually using an epithelial voltohmmeter 2 which was connected to a STX2 chopstick electrode (World Precision Instruments, USA). This was done to ensure stable TER. Organoid transwell monolayers were moved to a 24-well cell module cellZscope system (NanoAnalytics, Germany) with 1mL and 200µl of organoid media added to the basolateral and apical chambers respectively. This real-time recording system allows for parallel impedance spectroscopy measurements of monolayers grown on semi-permeable membranes. The cellZscope module, and organoid monolayers were housed in a humidified incubator (37L°C and 5% CO_2_) for the duration of experimentation with TER measurements taken every hour. Continuous TER measurements were recorded with raw TER data extracted and reported as TER measured over time. All experiments involving organoid monolayers were done using 6 biological replicates from each location (non-inflamed or prior inflamed) to reduce any monolayer-to-monolayer variability. Data was subsequently reported as an average for that location with each data point plotted representing those 6 monolayers per group. Analysis and calculations were performed in GraphPad Prism using ANOVA.

### Assay for Transposase-Accessible Chromatin sequencing (ATAC-seq) in organoids

Single cell suspensions of patient-derived organoids were prepared by trypsinization using TrypLE Express (Gibco, #12604013) until dissociation was achieved and followed by gentle mechanical disruption and strained with a 40 uM strainer (Falcon, #352340) and counted on the Thermo Fisher Countess II automated cell counter. ATAC-seq was performed according to Buenrostro et al.(38)and Corces et al.(39) Briefly, 50,000 cells were harvested and incubated at 4°C for 15 minutes in ATAC resuspension buffer [10mM Tris-HCl pH 7.4, 10mM NaCl, 3mM MgCl_2_, 0.1% NP-40, 0.1% Tween-20, 0.01% digitonin]. Nuclei were washed with the ATAC resuspension buffer without detergents followed by centrifugation at 500 x g for 10 minutes at 4°C. TDE1 enzyme (Illumina, cat no. 20034198) was used for tagmentation for 30 minutes at 37°C in 1X Tagment DNA Buffer, 0.02% Digitonin (Promega, cat no. G9441), 0.2% Tween-20. DNA was extracted using MinElute kit (Qiagen) and amplified using NEBNext High-Fidelity PCR Master Mix (NEB). DNA fragment selection was performed using sparQ PureMag Beads (Quanta bio, cat no. 95196). Quality of DNA was evaluated using the High Sensitivity D1000 ScreenTape (Agilent) run on a Tapestation 4150 (Agilent). Next-Generation sequencing was performed on NextSeq 2000 PE50 in the Genome Analysis Core in Mayo Clinic, Rochester. See Supplementary Methods for details on Bioinformatic Analysis for ATAC-seq.

### RNA sequencing in organoids

The same cells from organoids prepared for ATAC-seq were harvested by centrifugation and the pellet used for RNA extraction. RNA was extracted by miRNeasy Micro Kit (Qiagen) according to manufacturer’s instructions. RNA from treated organoids was extracted using miRNeasy Micro Kit (Qiagen) according to manufacturer’s instructions. Libraries were prepared using the stranded mRNA prep, ligation kit (Illumina) following the manual instructions. Quality of libraries was validated using Tapestation 4150 (Agilent). Next-Generation sequencing was performed on NextSeq 2000 PE50 in the Genome Analysis Core in Mayo Clinic, Rochester. See Supplementary Methods for details on Bioinformatic Analysis for RNA-seq.

### Organoid Exposure to Inflammatory Stimuli

Organoids were seeded at equal cell densities and treated with TNFα (Gibco, #AF-300-01A), which was supplemented in our 50% WRN media at a final concentration of 10 and 100 ng/ml for 24 hours. Organoids were harvested by removing TNFα supplemented media and adding 500 ul of Cell Recovery Solution to each dome (Corning, #354253). Organoids were placed on ice for 30 minutes in Cell Recovery Solution to remove Matrigel. Organoids were then washed 2x with PBS, and RNA was extracted and RNA-seq performed as described above.

### Organoid Wound Healing Assay

Single cells dissociated from intestinal organoids were seeded onto IBIDI 24-well culture inserts (IBIDI Cat.No: 80242) pre-coated with 2% Matrigel. Monolayers were allowed to proliferate until they reached a confluency of approximately 90-95%, at which point the insert was removed, creating two distinct wounds for each patient line. Images were obtained utilizing EVOS XL core microscope at a magnification of 4X over a duration of 3 days with designated time points captured at 0, 8, 20, 30, 48, 56 and 72 hours. Wound gaps were quantified using ImageJ, where normalized measurements were calculated to represent the percentage of wound area that was healed, with the initial time point (0 hours) serving as a reference point for 100% wound area. In total, four wounds from each patient line were evaluated and subjected to a two-way ANOVA followed by Sidak’s multiple comparisons test.

### Statistics

Statistical analyses were performed using GraphPad Prism 8.0.1 (GraphPad Software, Inc., San Diego, CA, USA), and R (4.0.3). For RNA-seq and ATAC-seq data, differential analysis was performed using DESeq2 and DiffBind, respectively, with multiple hypothesis testing correction applied (Benjamini-Hochberg FDR < 0.05), more details can be found in the Supplemental Methods section. The number of biological replicates and specific statistical tests used are indicated in figure legends.

### Study approval

This study was approved by the Mayo Clinic Institutional Review Board (IRB #21-006244). All human participants provided written informed consent prior to enrollment and tissue collection. All organoid derivation and experimental procedures were conducted in accordance with approved IRB protocols and institutional guidelines. No animal studies were performed.

### Data Availability

All sequencing data has been deposited at the Gene Expression Omnibus (GEO) with accession numbers GSE282442 (ATAC-seq) and GSE282444 (RNA-seq).

## RESULTS

### Inflammation alters the phenotype of Intestinal Stem Cells in human organoids

To examine how prior inflammatory exposure influences intestinal stem cell (ISC) behavior, we established paired human colonic organoids from inflamed (prior-inflamed, PI) and uninflamed (non-inflamed, NI) regions of the same ulcerative colitis (UC) patients (Figure 1A, Supplementary Table 1). We refer to organoids from inflamed tissue as “prior-inflamed” because they were isolated from sites of inflammation but subsequently cultured in the absence of inflammatory cues. While previous studies have successfully derived organoids from IBD patients, only a few have generated organoids from actively inflamed tissue, and none have performed matched comparisons of inflamed and uninflamed regions within the same individual.(35–37) This model enables direct intra-patient comparison while controlling for genetic and environmental variation, allowing us to assess persistent, epithelial-intrinsic changes in ISC function following inflammation. Once established *in vitro*, all organoids were maintained under identical, ISC-preserving conditions and passaged at least 10 times to eliminate residual immune and stromal influences, ensuring that any lasting molecular or functional differences reflected stable epithelial changes rather than transient inflammatory effects.

**Figure 1.**
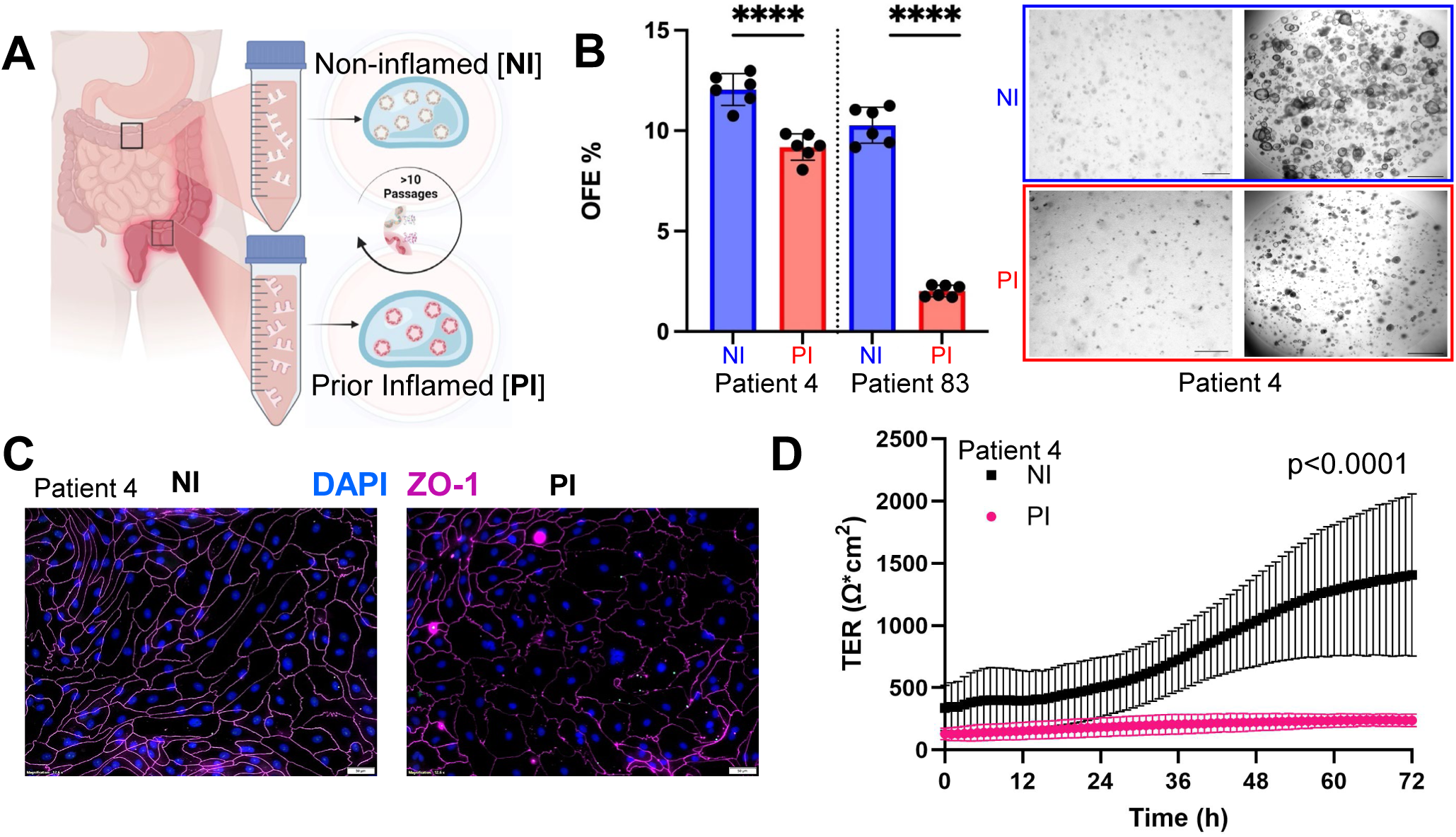
Human ISC-derived organoids from inflamed tissue are altered compared to matched organoids from uninflamed sites. (A) Schematic of paired sampling strategy showing organoid generation from inflamed (PI) and non-inflamed (NI) regions of the colon from the same patient with IBD. Organoids are cultured under identical conditions and are passaged for greater than 10 passages before any molecular profiling. (B) Organoid-forming efficiency (OFE) is significantly reduced in PI compared to NI organoids. A single cell suspension (left brightfield images) from dissociated organoids were plated identically between NI (top brightfield images) and PI (bottom brightfield images) and allowed to form organoids over 7 days (right brightfield images) for two patients, and quantified in the bar chart (on right). (C) Immunofluorescence staining of ZO-1 highlights disrupted tight junctions in PI monolayers. (D) Transepithelial electrical resistance (TER) is significantly decreased in PI (pink) epithelial monolayers compared to NI (black), over three days.

The microscopic appearance of the organoid cultures did not differ between NI and PI organoids, and the organoids maintained a cystic structure typical of colonic organoids (Figure S1A).(37) To evaluate the growth and regenerative capacity of ISCs from inflamed and non-inflamed regions, we measured organoid-forming efficiency (OFE), the ability of single cells to generate organoids under standardized culture conditions.(40) OFE serves as a functional readout of stem cell fitness and regenerative potential. Despite similar overall morphology in established cultures, we found that PI organoids exhibited a significantly reduced OFE compared to NI organoids (Figure 1B). We and others have reported disruptions in barrier proteins in IBD following inflammation,(41–43) and indeed the PI organoids showed that the tight junction protein ZO-1 was disrupted compared to NI organoids (Figure 1C and Figure S1B). This disruption in ZO-1 was confirmed as functional by measuring lower transepithelial resistance (TER) in the PI organoids compared to NI (Figure 1D), indicating barrier disruption. Together, these findings demonstrate that ISCs retain functional hallmarks of prior inflammation, even after prolonged culture in the absence of inflammatory cues.

### Intestinal Stem Cells retain an epigenetic signature of prior inflammation

The sustained epithelial dysfunction observed in PI organoids, even after removal from the inflammatory environment, prompted us to hypothesize that prior inflammation may reprogram ISC chromatin, similar to what has been reported in epidermal stem cells following inflammation.(21, 22) To investigate this, we performed assay for transposase-accessible chromatin using sequencing (ATAC-seq) to globally profile chromatin accessibility in ISCs preserved in PI and NI organoids (Figure 2A). This approach provides an epigenetically unbiased view of the genome-wide accessible chromatin landscape, enabling the identification of regulatory regions altered by prior inflammation without requiring prior knowledge of specific epigenetic marks or regulatory elements.

**Figure 2.**
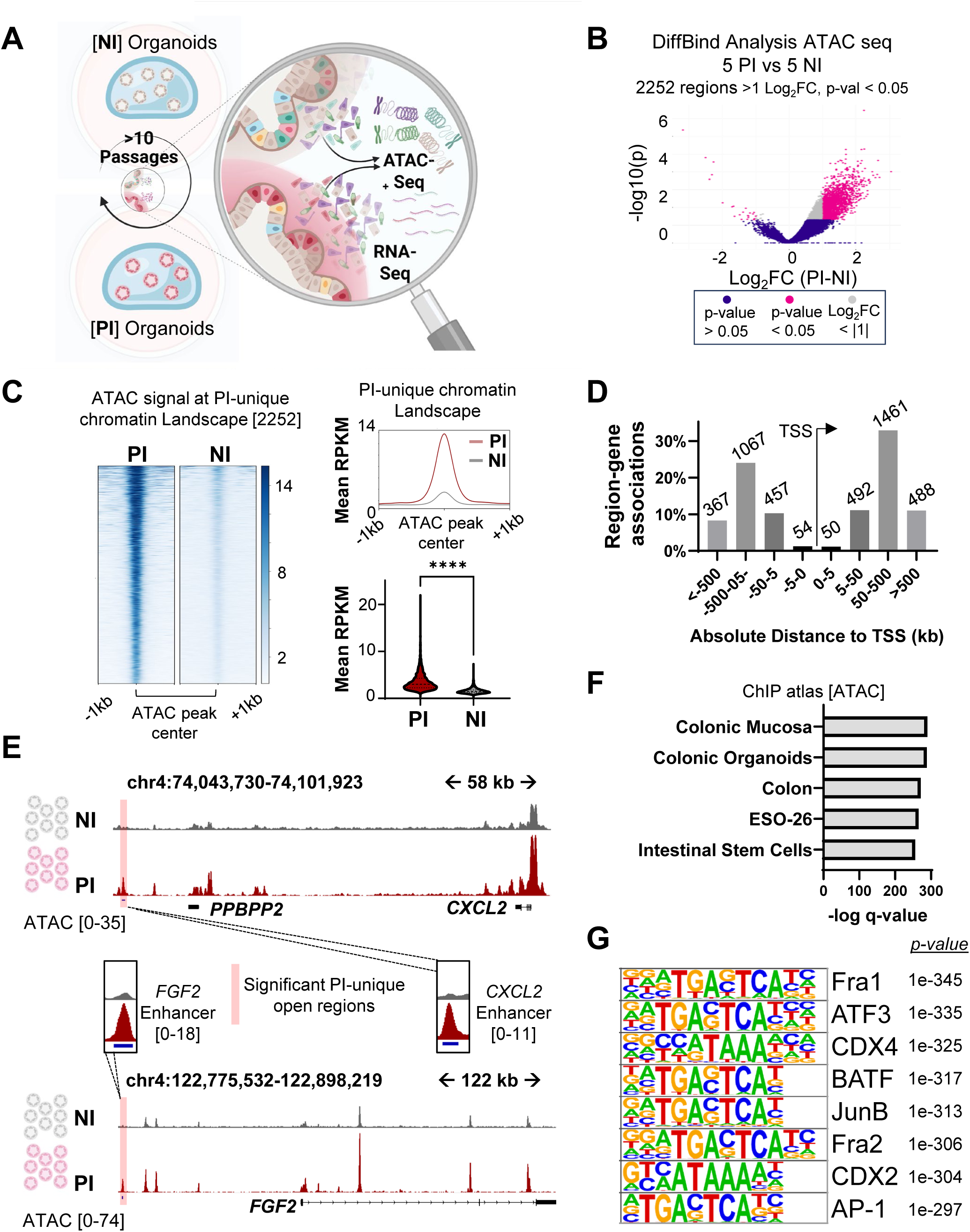
Chromatin Accessibility Is Altered in ISCs From PI Organoids. (A) Schematic of sequencing performed in NI and PI organoids; these are the identical organoids used throughout. (B) Volcano plot showing 2,252 PI-unique accessible regions (pink) consistently across multiple patients following analysis of differential chromatin accessibility between NI and PI organoids. (C) Average ATAC-seq signal at PI-unique peaks demonstrates significantly sustained accessibility in PI compared to NI. (D) Genomic annotation of open chromatin regions. (E) Genome tracks showing increased accessibility at CXCL2 and FGF2 loci in PI organoids. (F) Motif footprinting shows enrichment of AP-1 and CDX2 in PI-unique peaks. (G) Transcription factor motif enrichment analysis highlights AP-1 family members (JUNB, ATF3) and lineage-specific factors (CDX2) in PI-unique regions.

Analysis of differential chromatin accessibility revealed 2252 chromatin regions (PI-unique regions) that were more open in PI relative to NI, and that were consistent across all patients (Figure 2B). Correlation analysis showed strong concordance between duplicates and a higher agreement between patients over inflammation status (Figure S2A), while principal component analysis demonstrated a clear separation between PI and NI samples (Figure S2B). When examining the PI-unique regions, there was a significantly higher signal in PI compared to NI organoids, whether plotted together as consensus peaks for all patients or individually (Figure 2C; Figure S2C). To address the reproducibility and fundamental nature of our findings, we examined ATAC-seq signal in three additional UC patients (Patients 8, 38, and 129). Using the consensus set of PI-unique regions derived from five matched PI–NI pairs, we observed a similar enrichment of chromatin accessibility in these independent PI samples (Figure S2D).

The PI-unique chromatin regions were enriched primarily at enhancer regions (Figure 2D), which is reflected in the representative genome tracks showing sustained increased accessibility at enhancer regions at the CXCL2 and FGF2 loci in PI organoids (Figure 2E). We also observed similar and unchanged chromatin accessibility in NI-PI organoids pairs at EPCAM, a gene essential for epithelial identity and integrity, illustrating that key regulatory regions required for core epithelial function remain unchanged despite prior inflammatory exposure (Figure S3A). Overlapping our set of PI-unique regions with the ChIP atlas database(44) specific to publically available ATAC-seq data revealed the specificity of our system, including intestinal stem cells and colonic organoids (Figure 2F). To identify regulatory factors that may contribute to or result from persistent chromatin accessibility in PI ISCs, we performed motif enrichment analysis,(45) motif footprinting(46) and co-localization enrichment(47, 48) at the PI-unique open chromatin regions, which revealed strong enrichment for AP-1 factors (JUNB, ATF3), along with lineage specific binding motifs (CDX2), which was consistent using multiple different programs (Figure 2G, Figure S3B,C). The motifs enriched in NI organoids suggest that non-inflamed ISCs maintain transcriptional programs associated with homeostasis, epithelial barrier maintenance, and metabolic stability, with reduced accessibility in PI organoids perhaps reflecting a disruption of normal epithelial identity and function. These results demonstrate that prior inflammation leaves a lasting chromatin imprint in ISCs, characterized by sustained accessibility at enhancer regions and enrichment of AP-1 and intestinal lineage transcription factor motifs, supporting the existence of an epigenetic memory of intestinal inflammation.

### Inflammation-associated open chromatin reflects a transcriptionally primed state

To assess whether the persistent open chromatin regions identified in PI organoids were linked to transcriptional changes, we performed bulk RNA-seq on the same PI and NI organoids. To integrate this data with our ATAC-seq results, we mapped the 2,252 PI-unique open chromatin regions to their associated genes, (49) which resulted in 1,904 protein coding genes associated with the PI-unique chromatin landscape. We then examined transcriptional differences between PI and NI organoids across all patients. Despite marked differences in chromatin accessibility, few of the genes associated with accessible chromatin were differentially expressed between PI and NI organoids (Figure 3A). Only 4.8% of genes were significantly upregulated by differential expression analysis in all patients (fold change =>1, p-adjusted <0.05). This lack of transcriptional activation despite increased chromatin accessibility suggests that the regions are in an inactive, but primed regulatory state. Pathway enrichment analysis of genes linked to PI-unique chromatin regions revealed significant overrepresentation of biological programs associated with epithelial stress responses, regeneration, and inflammation, including MAPK, WNT, TGFβ signaling, and cellular senescence (Figure 3B).

**Figure 3.**
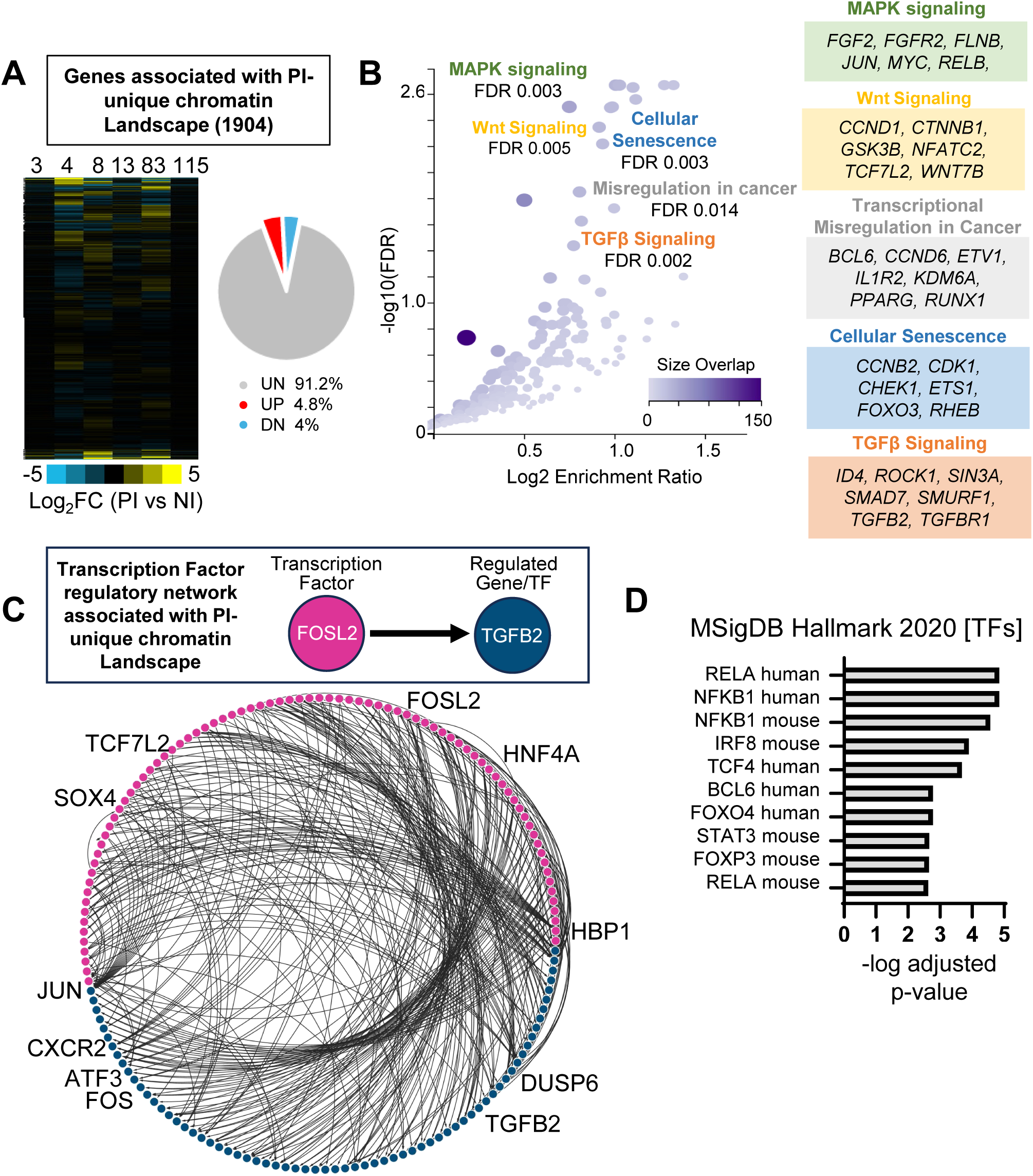
Genes Linked to PI-Unique Chromatin Regions Reflect Modest Transcription Activation. (A) Heatmap of Z-score FPKM expression values for 1,904 genes associated with PI-unique chromatin regions across six patients (left). Pie chart illustrating the percentage of genes that were unchanged (UN), upregulated (UP), or downregulated (DN) when comparing PI to NI at the 1904 genes. (B) Pathway enrichment of PI-associated genes using KEGG. (C) Transcription factor regulatory network reveals connectivity among TFs and their associated genes. Top box shows an example of iRegulon approach to determining regulation of TF with an associated gene. (D) Comparison of PI-unique regions specifically with transcription factor target gene enrichment from MSigDB.

To further investigate the transcriptional regulators driving these programs, we performed a transcription factor regulatory network analysis, which maps transcription factors to their putative target genes based on motif enrichment and cis-regulatory sequence conservation.(50) Among the genes associated with PI-unique accessible chromatin regions, ∼17% (322 genes) encoded transcription factors, including FOS, JUN, ATF3, TCF7L2, SOX4, and TGFB2, known regulators of inflammatory signaling and stem cell regulation (Figure 3C and S4A). We next cross-referenced PI-enriched regions with curated transcription factor target gene sets from the MSigDB Hallmark collection and found significant enrichment for targets of inflammatory transcription factors such as RELA (NF-κB), NFKB1, IRF8, STAT3, and TCF4 (Figure 3D), reinforcing the idea that inflammation leaves ISCs epigenetically primed at key regulatory hubs.

To this point, our analysis focused on genes linked to PI-unique open chromatin, revealing a regulatory landscape that is epigenetically accessible but largely transcriptionally inactive. To better understand the broader transcriptional shifts that may contribute to the PI phenotype, we next examined genome-wide expression differences between PI and NI organoids. Pairwise correlation analysis of gene expression showed that organoids from the same patient (PI and NI) were more similar to each other than to organoids from other patients, regardless of inflammation status, indicating patient-specific transcriptional identity is maintained (Figure S4B). Genes upregulated in PI organoids included IGFBP1, PRAC1/2, and BMP6, factors associated with stress responses, epithelial remodeling, and altered signal transduction, and downregulated genes such as PITX2, GATA5, and HOXC10, are developmental transcription factors often associated with tissue specification, differentiation, and epithelial identity (Figure S4C). Gene set enrichment analysis (GSEA) on the full transcriptome showed that PI organoids were enriched for genes downregulated by IL-6 deprivation, EGFR signaling, glycolysis, and proliferative cycling (Figure S4D), further supporting a shift toward a quiescent or stress-adapted transcriptional state.

Our findings reveal two distinct layers of transcriptional regulation in inflammation-experienced ISCs. Whole transcriptomic analysis revealed PI organoids exhibited elevated stress- and epithelial remodeling-associated programs and reduced expression of differentiation factors, features that may underlie the impaired regeneration and barrier dysfunction. In parallel, the genes associated with PI-unique open chromatin regions were not transcriptionally upregulated. Given their enhancer-like features and enrichment for key transcription factor motifs, we hypothesized that these regions may represent regulatory elements primed for rapid activation upon challenge. Thus, we next asked whether PI-unique accessible genes mediate a heightened transcriptional response following inflammatory stimulus.

### Prior inflammation enhances ISC response to secondary inflammatory challenge

The open chromatin regions identified in PI organoids raised the possibility that prior inflammation had imprinted an epigenetic memory of a primed transcriptional state in ISCs. We hypothesized that these open chromatin regions function to endow the cell with the ability to respond rapidly to a secondary inflammatory challenge.

To functionally test this, we re-challenged matched PI and NI organoids with tumor necrosis factor alpha (TNFα) and performed two sets of RNA-seq experiments to evaluate gene expression changes (Figure 4A). TNFα was selected as the inflammatory stimulus given that TNFα is a well-characterized upstream activator of AP-1 and has been previously used to model inflammatory signaling in intestinal organoids derived from IBD patients. (51) We performed a dose-response RNA-seq experiment in organoids from Patient 4, treating NI and PI organoids with 10 ng/µL and 100 ng/µL TNFα, which revealed a broader transcriptional response in PI organoids across both concentrations (Figure S5A–B). Based on these findings, we proceeded with RNA-seq in organoids from two patients (Patients 4 and 83) treated with 10Lng/µL TNFα.

**Figure 4.**
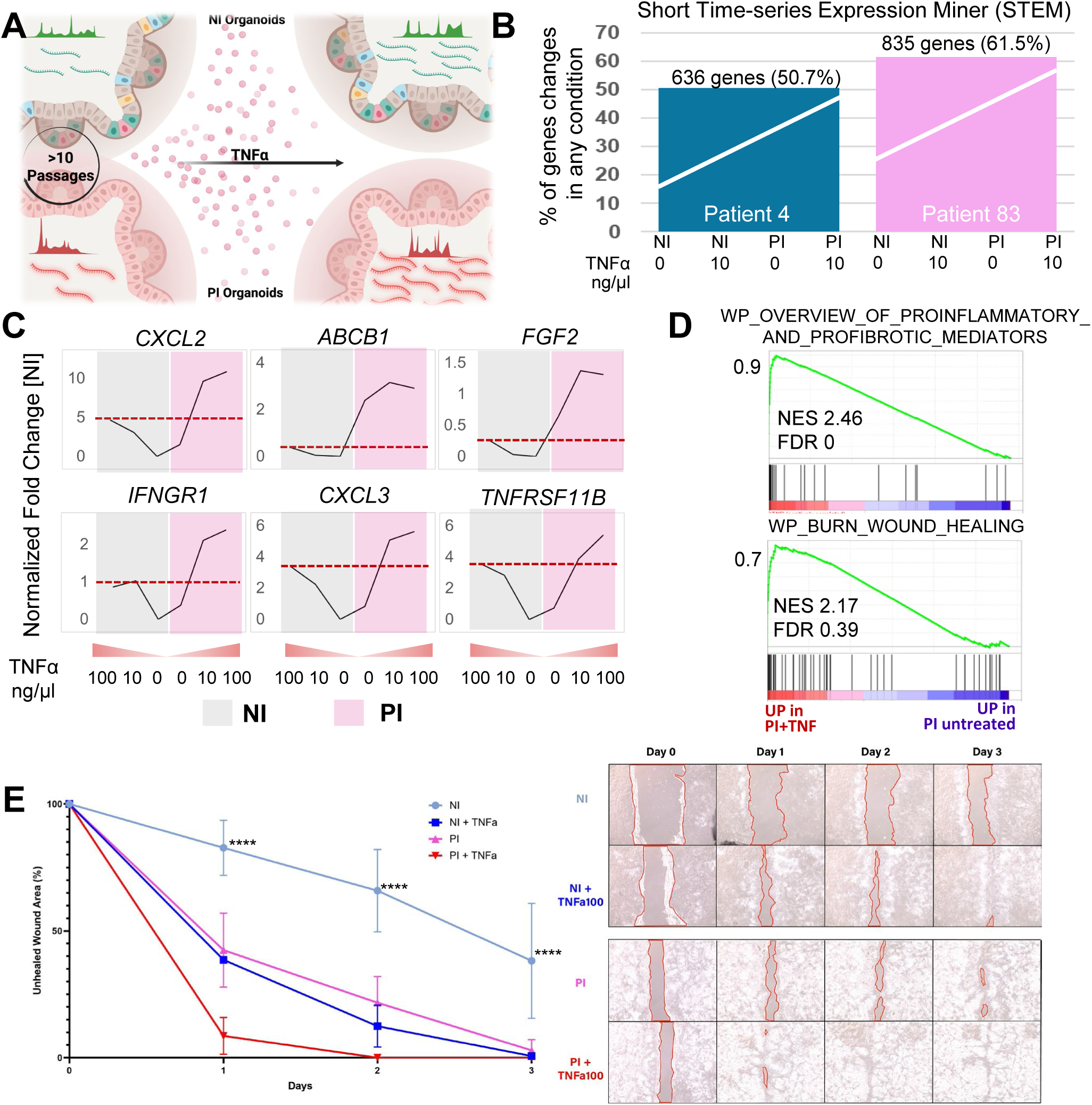
PI Organoids Exhibit Enhanced Transcriptional Responses to Inflammatory Re-challenge. (A) Experimental design of TNFα re-stimulation in NI and PI organoids. (B) STEM analysis indicates a higher percentage of transcriptionally responsive genes in PI vs. NI organoids. (C) Expression profiles of representative TNFα-responsive genes across increasing TNFα concentrations (0, 10, 100 ng/μL) in PI (pink) and NI (gray) organoids. Genes such as CXCL2 and CXCL3 showed enhanced induction in PI organoids. (D) Gene set enrichment analysis revealed increased activation of pathways related to proinflammatory mediators (top) and wound healing (bottom) in PI organoids after TNFα treatment. (E) Wound healing assay in organoid-derived monolayers. **** p-value<0.0001, and represents all comparisons to NI organoids, which remained significant throughout conditions and time points. Left: Quantification of wound closure over 3 days. Right: Representative images of wound closure across time points.

To assess dominant transcriptional responses to TNFα in NI and PI organoids, we clustered differentially expressed genes based on shared expression trajectories using Short Time-series Expression Miner (STEM).(52) In both patients, the most significantly enriched cluster represented genes more strongly induced in PI organoids following TNFα treatment, comprising 50.7% and 61.5% of PI-unique genes with open chromatin, indicating a broader transcriptional response in inflammation-experienced epithelium (Figure 4B and Supplementary Figure S6A and B). Analysis of representative PI-unique genes with open chromatin revealed more robust upregulation in the PI organoids compared to NI in both experiments (Figure 4C; Figure S5C). GSEA further demonstrated that the genes that were more robustly reactivated in the PI organoids were enriched for proinflammatory and profibrotic gene programs and pathways linked to wound healing (Figure 4D). To evaluate whether this transcriptional memory impacted epithelial function, we performed a scratch assay in monolayers derived from NI and PI organoids, with or without TNFα exposure. PI organoids exhibited significantly enhanced closure capacity compared to NI organoids at each time point across the duration of 3 days (Figure 4E). Upon TNFα exposure, both NI and PI organoids showed improved closure, but the response was significantly greater in PI organoids, suggesting that inflammation-experienced ISCs are functionally primed to mount an amplified regenerative response upon secondary inflammatory challenge. Taken together, these findings indicate that ISCs do not return to a naïve state after inflammation but instead retain a molecular memory that shapes future cellular responses to inflammatory injury.

## DISCUSSION

This study uncovers a previously unrecognized layer of stem cell regulation in IBD, by providing direct evidence that the intestinal epithelium retains a durable epigenetic memory of prior inflammation. While prior studies have shown that IBD-derived organoids retain features of epithelial dysfunction,(31–34) none have interrogated chromatin alterations in matched inflamed and uninflamed regions from the same patient.(35–37) Using paired human colonic organoids derived from inflamed and uninflamed regions of the same UC patients, we demonstrate that prior inflammatory exposure leaves a stable chromatin imprint in ISCs, characterized by sustained accessibility at enhancer-like regulatory elements, enrichment for AP-1 transcription factor motifs, and enhanced transcriptional responsiveness to subsequent inflammatory stimuli (Figure 5). Because organoid cultures are initiated and maintained by ISCs, persistent epigenetic features must reflect stem cell-intrinsic memory mechanisms. These findings establish that ISCs are not merely reset to a naïve state following resolution of inflammation but instead maintain a molecular memory that can impact future cellular behavior. Future work using single-cell chromatin and transcriptomic profiling will be necessary to confirm ISC-specific memory signatures and determine how these epigenetic programs are propagated into progeny lineages during epithelial renewal.

**Figure 5.**
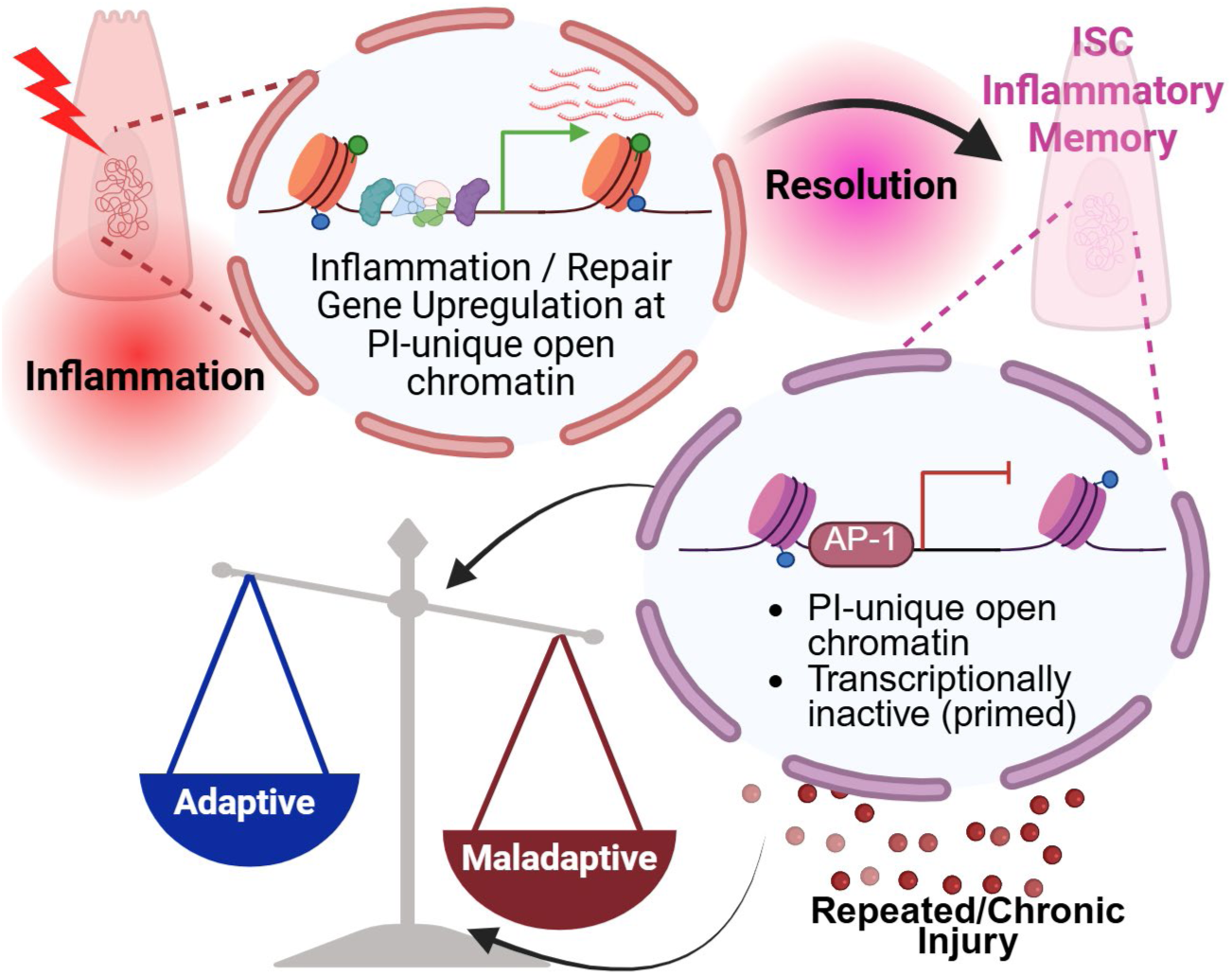
Model of Inflammatory Memory in Intestinal Stem Cells. Inflammation induces chromatin remodeling in ISCs at enhancer regions linked to genes involved in inflammation and tissue repair. After inflammation resolves, a subset of regions remains open, establishing a primed chromatin state characteristic of inflammatory memory. This memory endows ISCs with enhanced transcriptional responsiveness upon secondary inflammatory challenge. While this primed state can support adaptive responses, such as rapid reactivation of gene programs and healing following re-challenge, it may also predispose to maladaptive outcomes under conditions of repeated or chronic injury.

The persistent chromatin accessibility observed in inflammation-experienced organoids was concentrated at enhancer-like regions near genes involved in epithelial stress responses, regeneration, and inflammatory signaling. Although many of these regions were transcriptionally silent under homeostatic conditions, challenge with TNFα showed enhanced transcriptional activation in inflammation-experienced organoids (PI) compared to uninflamed organoids (NI).

These data support the interpretation that inflammation-experienced ISCs exist in a transcriptionally primed state defined as silent under homeostasis but rapidly inducible upon secondary challenge.

This memory may serve an adaptive function, enabling ISCs to respond more efficiently to recurrent insults and support rapid epithelial regeneration. Indeed, inflammation-experienced organoids displayed enhanced wound healing responses compared to uninflamed organoids at baseline and when re-exposed to inflammatory stimuli, suggesting that prior inflammatory exposure may prime ISCs for accelerated repair. However, our findings also raise the possibility that this same epigenetic imprinting may become maladaptive, predisposing the epithelium to dysregulated repair, persistent activation of inflammatory pathways, or eventual progression to neoplasia. In the context of UC, where disease recurrence often arises at previously affected sites despite histologic healing,(53) such persistent reprogramming of ISCs may contribute to the chronic and relapsing nature of the disease. Whether inflammatory memory in ISCs influences disease recurrence remains an open and critical question.

This work highlights the power of human organoid systems for uncovering stable, cell-intrinsic features of disease that are not captured by short-term transcriptomic profiling alone. While further studies are needed to define how inflammatory memory evolves over time and whether it can be erased or reprogrammed, our findings provide a foundation for understanding how ISCs integrate past inflammatory experience into future behavior. Targeting memory-associated transcription factors or chromatin remodelers may offer novel therapeutic strategies to modulate epithelial responses and promote mucosal healing in patients with chronic disease. This opens the door to new therapeutic strategies aimed at modulating the epigenetic landscape of ISCs to either enhance regenerative responses in acute inflammation or dampen maladaptive programs in chronic disease. Understanding how to balance the protective versus pathogenic aspects of inflammatory memory will be critical for designing interventions that promote long-term mucosal healing without fueling inflammation-associated complications.

In summary, we demonstrate that that inflammation leaves a lasting molecular imprint in ISCs from UC patients that shapes their future responses to injury. This memory is encoded in accessible chromatin landscapes that potentially primes cells for enhanced transcriptional and functional responses to subsequent insult. These findings provide a foundational framework for understanding how stem cells integrate past inflammatory exposures into future tissue behavior, an insight that may help reframe epithelial resilience and vulnerability in chronic diseases such as IBD. Inflammatory memory in the intestinal epithelium may represent a double-edged sword: supporting regenerative readiness on one hand, while perpetuating disease progression on the other. Understanding and modulating this memory may open new avenues for promoting durable healing and preventing relapses in patients with IBD.

## Author contributions

F.H.H.: investigation, analysis and interpretation of data, statistical analysis, and writing, review & editing; W.A.F: study concept and design, funding acquisition, writing, review & editing; B.R.D.: study concept and design, project supervision, funding acquisition, investigation, analysis and interpretation of data, drafting of the manuscript, writing, review & editing; all other authors: investigation, review & editing.

---

**FigS1.**
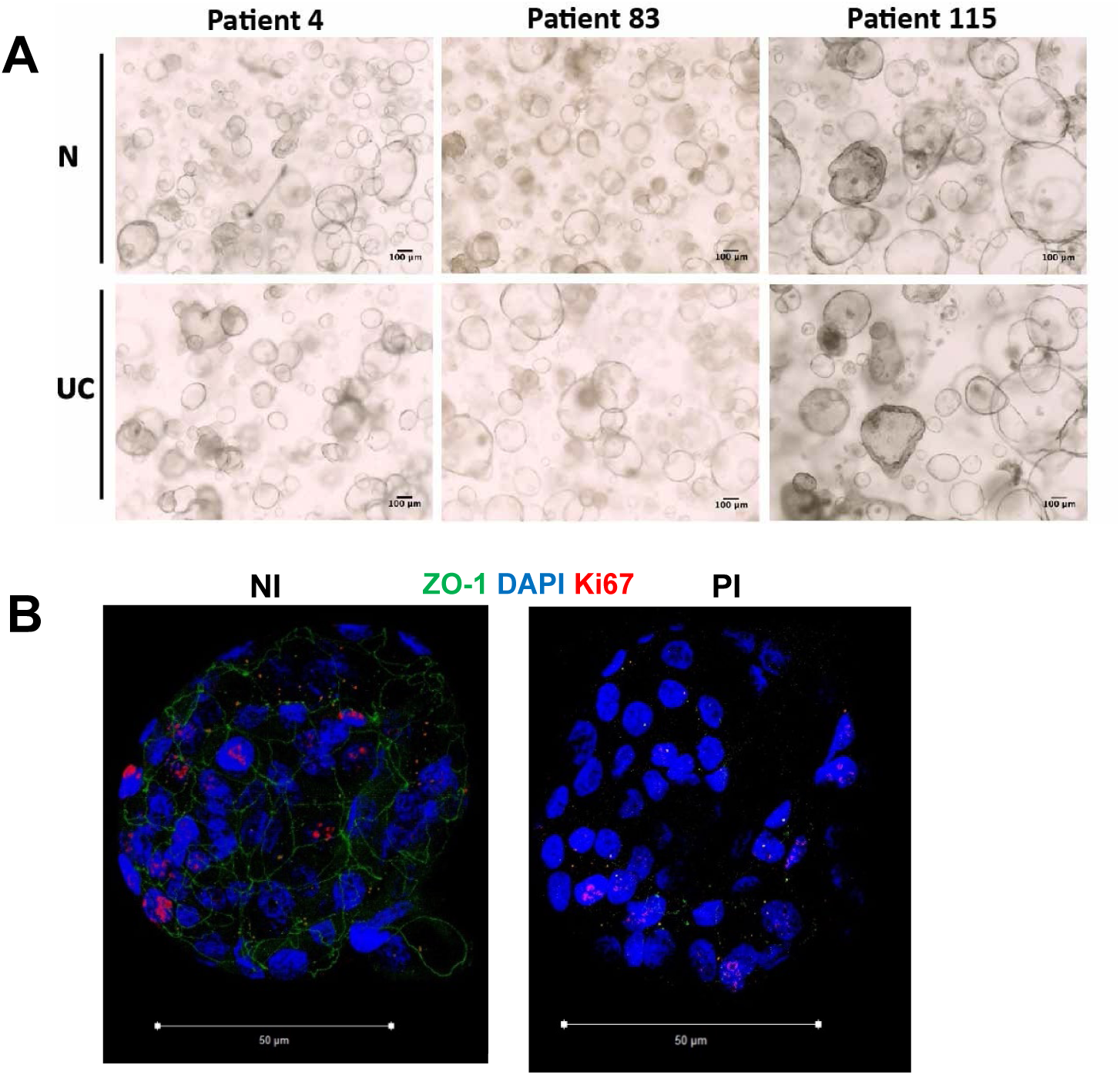

**FigS2.**
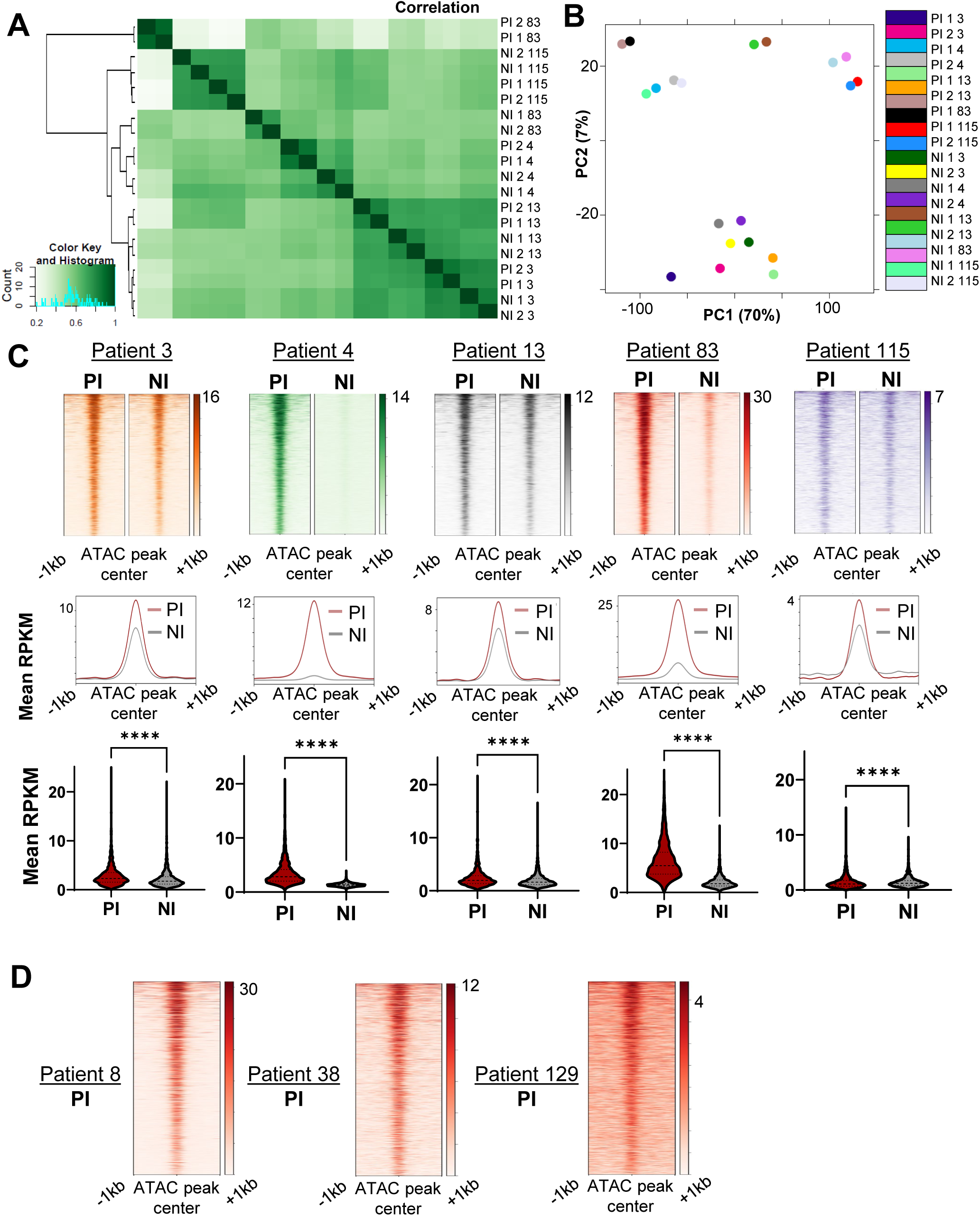

**FigS3.**
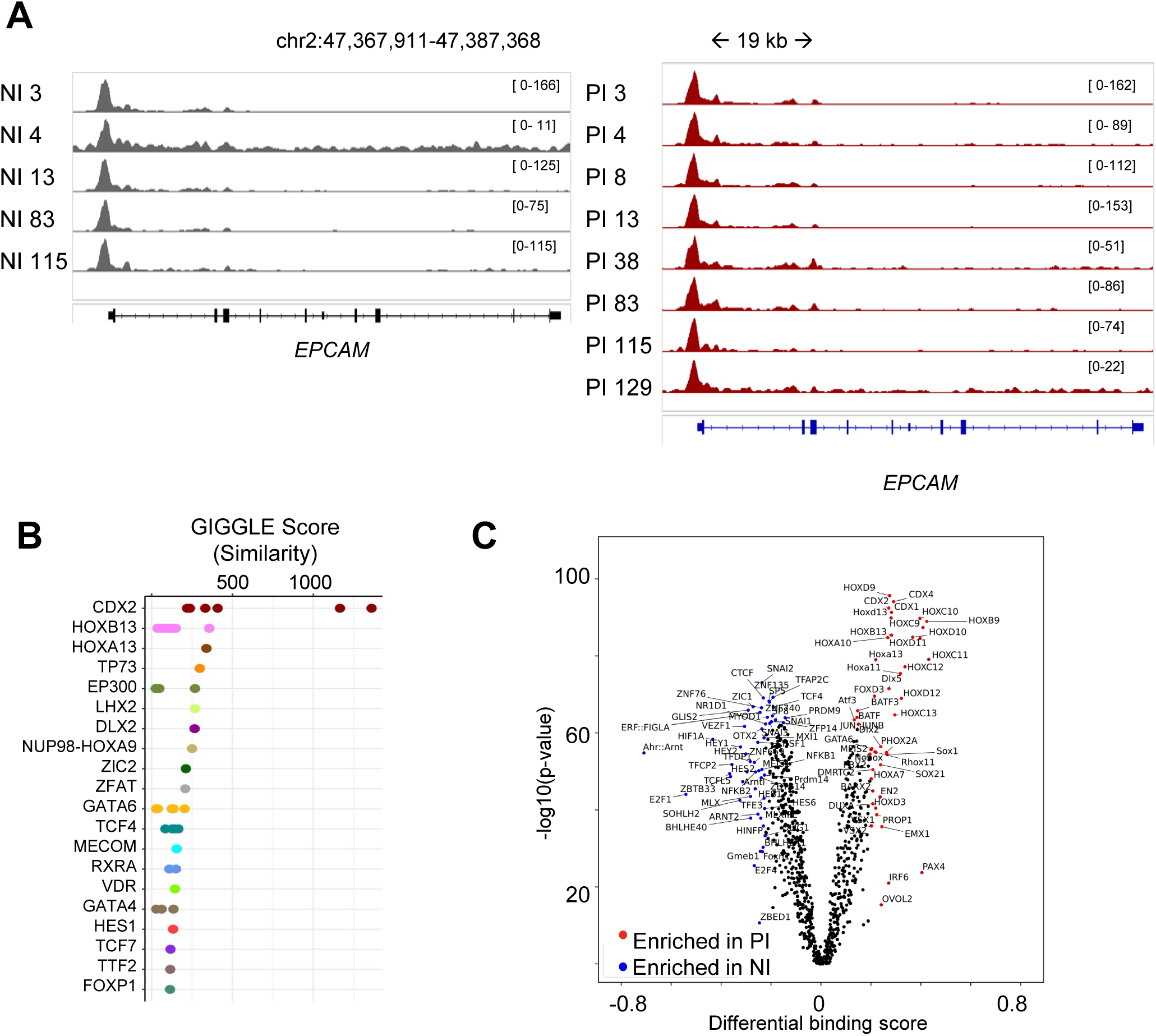

**FigS4.**
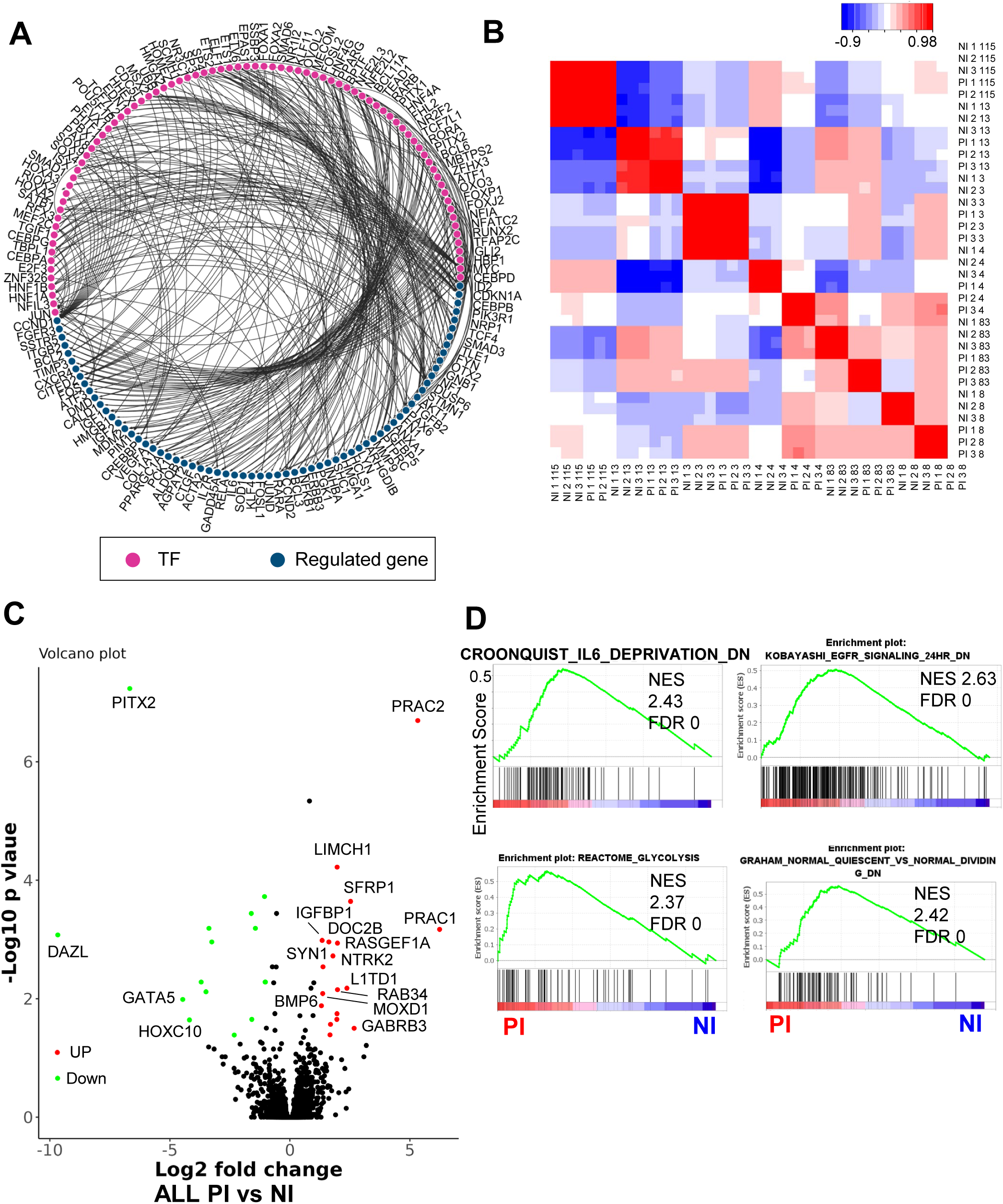

**FigS5.**
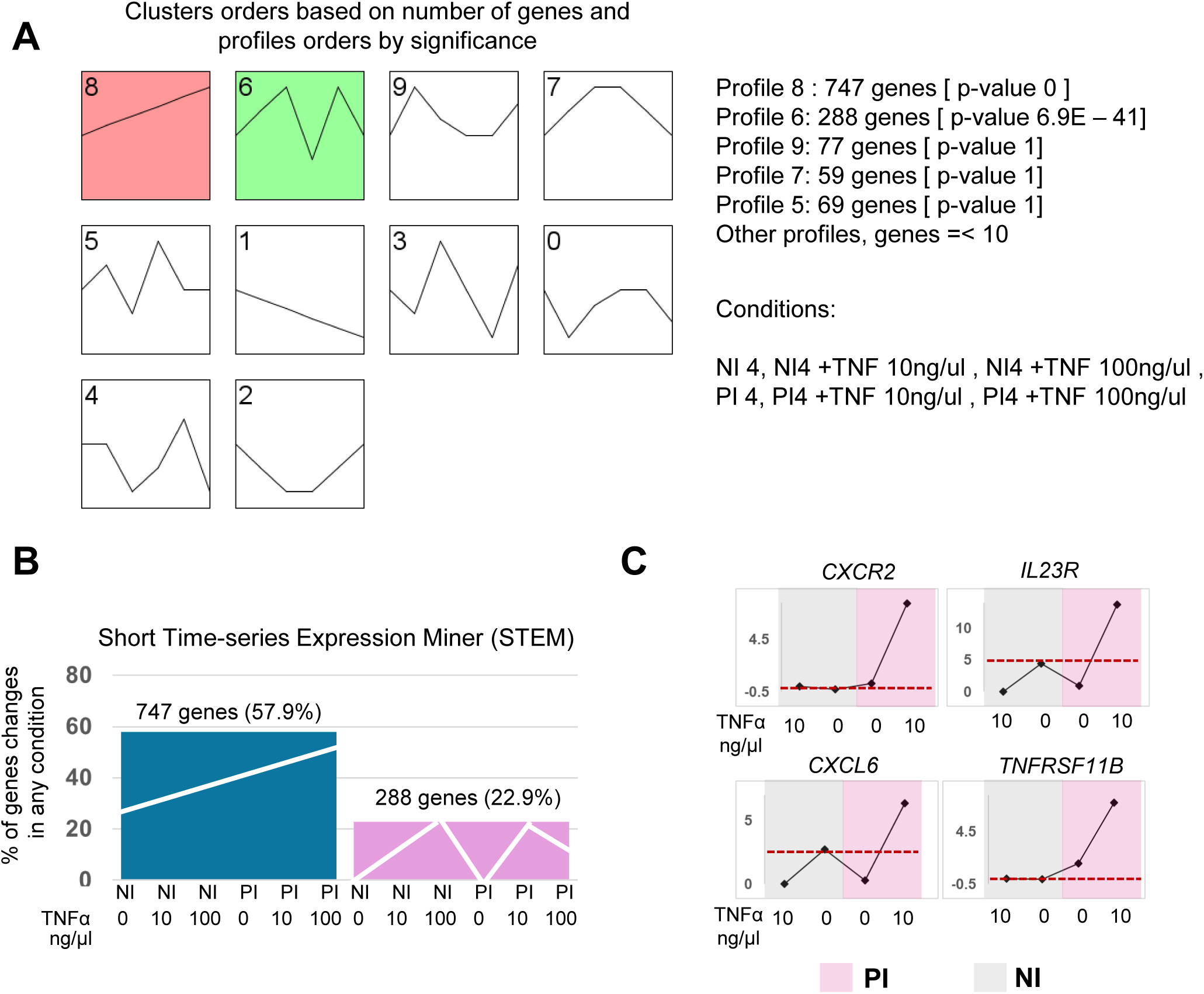

**FigS6.**
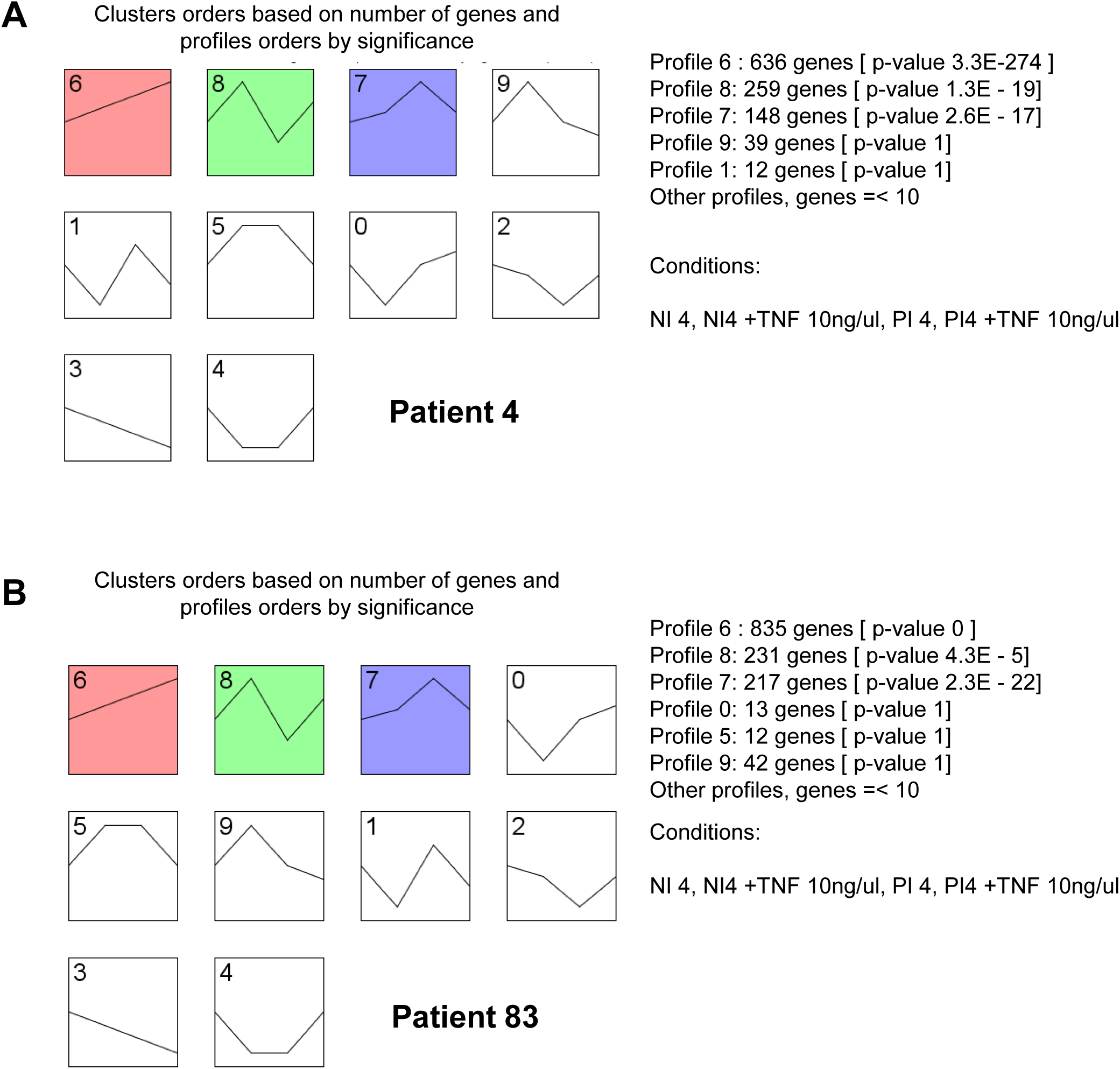

