## Supplemental Material for "Intestinal Stem Cells from Patients with Inflammatory Bowel Disease Retain an Epigenetic Memory of Inflammation"

**Supplementary Methods**

Human Colon Organoid Media Composition

Human Colon Media [ADMEM containing 50% Wnt, R-Spondin, and Noggin (WRN) from conditioned media of L-WRN cell line (ATCC)] and supplemented with: 1x N2 supplement, 1x B27 supplement, 40 ng/mL EGF, 3 uM SB202190, 500 nM A 83-01, 10 uM Y-27632, 1 uM NAC, 10 mM Nicotinamide, 10 nM Gastin I, 100 ug/mL Primocin, 1x Antibiotic/Anti-mycotic]

Seeding organoids as monolayers on transwell cell culture inserts

A 2% Matrigel solution was made by diluting Matrigel in PBS (Corning, #35623). 200 ul of the 2% Matrigel solution was placed on the upper (apical) reservoir in a 24 well plate with a transwell cell culture insert with a 3.0 uM pore size (Corning, #CLS3415) for 2-3 hours in an incubator at 37 °C. Organoids were dissociated into single cells using TrypLE Express (Gibco, #12604013) and were strained using a Falcon 40 uM cell strainer (Corning, #352240). The single cell suspension was counted on a hemocytometer. The 2% Matrigel solution was removed, and 3x10^5 cells of the single cell suspension were placed in the apical reservoir on the transwell insert. The total volume added to the apical reservoir with the single cell suspension was 200 ul, and 600 ul of our 50% WRN medium was added to the lower (basal) reservoir. Media was changed every 2-3 days until a monolayer covered the insert of the transwell.

Bioinformatic analysis in patient-derived organoids

Quality of reads from fastq raw files was evaluated using fastqc (v0.11.9) to check for base quality scores, adapter contamination, and sequence duplication levels. Reads passing quality metrics were mapped to hg38 (GRCh38, GCA_000001405.15) using bwa (v0.7.17)^1^ with parameters -n 10 -a 500 -o 10000 -N 10 -s. Only uniquely mapped, non-duplicated reads with mapping quality (MAPQ > 30) were retained using samtools (v1.18).^2^ Narrow peaks were called using MACS2 (v2.0.10)^3^ with parameters: -f BAM -g hs --keep-dup all -q 0.01. Peak lists were then filtered to remove blacklisted regions (hg38-blacklist.v2.bed). To compare and visualize peak intensities across samples and conditions, normalized bigwig (bw) files were generated using the bamCoverage tool in deeptools (v3.5.1).^4^ UC-specific regions were defined using the Bioconductor R package (Diffbind).^5^ Differential accessibility analysis between inflamed and uninflamed groups was conducted using DiffBind (v2) in R (v4.2.2), applying edgeR for statistical modeling using default parameters and minMembers=3. Differentially accessible regions with a pvalue < 0.05 and |log2 fold change| > 1 were retained for downstream analysis. Deeptools (v3.5.1) was used to generate heatmaps and binding profiles. Associated genes with regions were defined using GREAT analysis.^6^ Association rule used was Basal+extension: 5000 bp upstream, 1000 bp downstream, 5Mb max extension. Transcription factor motifs in the differentially accessible regions were identified using HOMER’s findMotifsGenome.pl utility using default parameters.^7^ To further analyze transcription factor binding at differentially open chromatin regions, TOBIAS (v0.13.3)^8^ was used for footprinting. BAM files from all replicates were merged for each condition before footprinting analysis. TOBIAS was run with default parameters for bias correction (ATACorrect), footprinting score calculation (ScoreBigwig), and transcription factor binding activity prediction (BINDetect). The JASPAR human core motif database was used as the motif set for TOBIAS BINDetect. Genes associated with differentially enriched peaks were analyzed for pathway enrichment using Enrichr web-based interface.^9–11^ To identify potential upstream regulators of differentially enriched genes, CistromeDB Toolkit^12,13^ and ChIP-Atlas^14^ were used. Gene lists derived from differential peak analysis were submitted to CistromeDB, which provided rankings of candidate transcriptional regulators based on their binding enrichment. Additionally, ChIP-Atlas was used to cross-reference gene targets with publicly available ChIP-Seq datasets, further confirming key regulators. Genome-wide signal tracks were visualized in IGV (v2.3.93).^15,16^ All analyses were conducted in a Linux-based computing environment running R (v4.2.2), Python (v3.10.7), and Java (JDK 13.0.2). Custom scripts for additional processing steps are available upon request.

Bioinformatic analysis for RNA-seq in patient-derived organoids

Raw paired-end sequencing reads were processed through the Mayo RNA-Seq bioinformatics pipeline, MAP-RSeq (v3.1.4).^17^ Briefly, Fastqc (v0.11.8) was used to assess read quality and Fastp (0.20.0)^18^ was used to trim adapter sequences. Trimmed reads were aligned to the hg38 reference genome using the splice-aware STAR aligner (v2.6.1c)^19^ using strand-specific basic two-pass mapping. To detect chimeric transcripts, chimeric reads with minimum segment length of 12 nucleotides were retained, and junctions with minimum overhang of 12 nucleotides were reported. Additionally, the minimum overhang for annotated splice junctions was set to 10 nucleotides. The maximum intron length and gap between read mates was set to 200 kb. Aligned and unmapped reads were reported in the resulting coordinate-sorted BAM file. In addition, MultiQC (v1.14)^20^ and RSEQC^21^ were used for comprehensive quality control of the aligned reads. Gene-level expression quantification was performed using Subread featureCounts (v1.6.3)^22^ with parameters -O -p -s 2 to obtain strand-specific raw gene counts and normalized (FPKM – Fragments Per Kilobase per Million mapped reads) counts. Pairwise differential expression analyses were conducted with raw gene-level counts using edgeR (v3.40.2)^23^ with default parameters. Genes were considered differentially expressed with Log2 Fold change >1 or < 1 and p-adjusted value < 0.05. To identify biologically enriched pathways, we used enrichR (v3.2)^9^ and Gene Set Enrichment Analysis (GSEA)^24^ got expressed genes using human gene set collections from the Molecular Signatures Database (MSigDB v2023.2.Hs) limiting to gene sets including 30-1000 genes.^25^ Differently expressed genes were hierarchically clustered using Cluster 3.0^26^ using Euclidean distance as a similarity metric and average linkage as a clustering method. Heatmaps were generated using Java Treeview 3.0.^27^ Pathway enrichment analysis was performed in WEB-based GEne SeT AnaLysis Toolkit (Webgestalt)^28,29^ using the following parameters: Over-Representation method, number analytes for category 30-2000, multiple test adjustment BH and a top 10 significance level. Differentially expressed transcription factors were used to build a TF-gene regulatory network using the TF-Target query of the iRegulon app^30^ in Cytoscape 3.10.3.^31^ To characterize gene signature behavior across TNF treatment, the short time-series expression miner (STEM)^32^ was used with the following parameters: normalize data, STEM Clustering method, maximum number of model profiles set to 10 and maximum unit change in model profiles between points set to 3. Graphs were designed using GraphPad Prism 8.0.1 (GraphPad Software, Inc., San Diego, CA, USA), BioRender.com and R (4.0.3).

**Supplementary Table 1.** Patient Clinical Data.

| **Sample ID** | **Sample Date** | **Mayo Score** | **Steroid?** | **Medications** | **Pathology Report** |
| --- | --- | --- | --- | --- | --- |
| IBD_83UC | 8/22/2022 | 2 | Yes | None | A. Colon and rectum, total proctocolectomy: Chronic ulcerative colitis, moderately active |
| IBD_83N | 8/22/2022 | 0 | Yes | None |  |
| IBD_13N | 7/28/2021 | 0 | No | Entyvio | D. Colon, sigmoid, endoscopic biopsy: Colonic mucosa with no diagnostic abnormality. No activity, granulomas, or dysplasia |
| IBD_13UC | 7/28/2021 | 1 | No | Entyvio | E. Colon, rectum, endoscopic biopsy: Colonic mucosa with no diagnostic abnormality. No activity, granulomas, or dysplasia |
| IBD_8UC | 6/2/2021 | 2 | Yes | Vedolizumab 600 mg every 4 weeks, infliximab 20 mg/kg every 4, azathioprine 125 mg daily, allopurinol 100 mg daily, Uceris 9 mg BID and topical and oral 5-ASA. | D. Colon, sigmoid, rectum, endoscopic biopsy: Malakoplakia. Separate fragments of colonic mucosa with Paneth cell metaplasia |
| IBD_3N | 4/26/2021 | 0 | No | Mesalamine | A. Colon, cecum, ascending, endoscopic biopsy: Minimal crypt architectural changes only. Consistent with inactive chronic colitis or other quiescent crypt destructive process. |
| IBD_3UC | 4/26/2021 | 2 | No | Mesalamine | D. Colon, sigmoid, rectum, endoscopic biopsy: Mild active chronic colitis without granulomas. No dysplasia |
| IBD_115N | 3/3/2023 | 0 | No | Ustekinumab; Imodium | Surgery tissue; He had inflammatory changes to his entire rectum and proximal sigmoid colon, distal sigmoid had uninflamed colon, cecal patch of inflammation was also removed. |
| IBD_115UC | 3/3/2023 | 3 | No | Ustekinumab; Imodium | A. Colon, subtotal colectomy: Active chronic colitis with mild to moderate activity, consistent with ulcerative colitis. No dysplasia. |
| IBD_4N | 5/3/2021 | 0 | Yes | Entyvio, Imuran, Prednisone 30mg |  |
| IBD_4UC | 5/3/2021 | 3 | Yes | Entyvio, Imuran, Prednisone 30mg | A. Colon, abdominal, and omentum, subtotal colectomy and omentectomy: Active chronic colitis with ulceration, consistent with ulcerative colitis . No dysplasia. |
| IBD_38UC | 2/15/2022 | 2 | No | Xeljanz | A. Colon and terminal ileum, subtotal colectomy: The colonic mucosa shows moderately to markedly active colitis with continuous involvement from the sigmoid colon proximally to the ascending colon, marked mucosal atrophy, scattered inflammatory pseudopolyps and diffuse thickening of the muscularis mucosa, consistent with the known history of ulcerative colitis. No dysplasia or granuloma seen. |
| IBD_129UC | 4/26/2023 | 1 | No | Entyvio | A. Colon, right hemicolectomy: Segment of colon with mild active inflammation. |

**SUPPLEMENTARY FIGURE LEGENDS**

**Figure S1. Phenotypic Differences Between PI and NI Organoids**
(A) Representative brightfield images of organoids derived from non-inflamed (NI) and inflamed (PI) regions of the colon from three UC patients. Scale bar = 100 μm. (B) Confocal images of organoids stained for ZO-1 (green), DAPI (blue), and Ki67 (red) show disrupted tight junctions and reduced proliferative cells in PI organoids compared to NI organoids. Scale bar = 50 μm.

**Figure S2. Chromatin Accessibility Patterns in Matched NI and PI Organoids**
(A) Sample-to-sample correlation heatmap showing high intra-patient similarity in ATAC-seq profiles across PI and NI organoids. (B) Principal component analysis of ATAC-seq data shows distinct separation of NI and PI samples along PC1. (C) Heatmaps and average signal plots of ATAC-seq signal centered on PI-unique peaks. Violin plots of mean RPKM values of ATAC peaks indicate statistically significant differences (****P < 0.0001). (D) Heatmaps show enrichment of PI-unique peaks in additional independent PI-only organoid samples (Patients 8, 38, and 129).

**Figure S3. PI Organoids Exhibit Conserved Epithelial Identity and Enrichment of Inflammatory and Lineage-Specific Motifs**
(A) Genome browser tracks show conserved accessibility at the epithelial marker EPCAM in both NI and PI organoids. (B) GIGGLE similarity analysis at PI-unique open chromatin. (C) Volcano plot of differential motif binding scores between PI and NI organoids highlights motifs enriched in PI (red) and NI (blue).

**Figure S4. Differential Transcriptional Profiles and Regulatory Network Analysis**

(A) TF-gene regulatory network from Figure 3C listed all TFs and regulated genes, only a subset are listed in the main figure. (B) Correlation matrix of RNA-seq profiles for each PI and NI samples, in triplicate. (C) Volcano plot of differentially expressed genes between all PI and NI organoids. Upregulated genes in red; downregulated genes in green. (D) GSEA plots for all RNA-seq data.

**Figure S5. TNF-Induced Transcriptional Reactivation in PI Organoids**
(A) STEM analysis of PI and NI organoids treated with 10 ng/uL or 100 ng/uL TNFα. Profile 8 (red highlighted) captures genes specifically reactivated in PI organoids. (B) Summary of STEM analysis shows that 747 genes (57.9%) show this pattern of enhanced expression in PI organoids treated with TNFα, while 22.9% of genes are reactivated by TNFα in both NI and PI. (C) Line plots of representative genes (CXCR2, IL23R, CXCL6, and TNFRSF11B.

**Figure S6. Consistent PI-Specific Transcriptional Reactivation in Two Patients**
(A–B) STEM clustering of gene expression changes in response to TNFα in NI and PI organoids from Patient 4 (A) and Patient 83 (B). Profile 6 shows consistent upregulation in PI organoids.
